# Dual Transcriptomic and Epigenomic Signatures of THC Exposure in Human Prefrontal Cortex Development

**DOI:** 10.64898/2026.01.08.698442

**Authors:** Luke Daniel L. Ofria, David W. Sosnowski, Aishwarya Pantula, Kaustubh Joshi, Brion S. Maher, Annie Kathuria

## Abstract

Prenatal cannabis use is increasing globally, with estimates up to 35% in North America. Fetal exposure to Δ9-tetrahydrocannabinol (THC) has been linked to neurodevelopmental deficits. Yet mechanistic understanding remains limited because animal models incompletely recapitulate human fetal development, and human-relevant in vitro platforms are scarce. To address this gap, we generated human iPSC-derived prefrontal cortex organoids (PFCOs) and used an integrated multi-omics approach combining bulk RNA-seq, whole-genome bisulfite sequencing (WGBS), and electrophysiology to characterize early molecular and functional responses to acute THC exposure at a developmentally relevant stage. THC induced a rapid, transient shift toward excitatory and neurodevelopmental gene expression programs while simultaneously suppressing extracellular matrix and adhesion pathways critical for structural support. Concurrent epigenetic remodeling selectively targeted synaptic assembly, postsynaptic organization, and axonal guidance genes, creating a mismatch between early activation of neuronal programs and epigenetic repression of the scaffolding required for their proper integration. Functionally, these disruptions manifested as delayed but reversible increases in burst duration at 24 hours, indicating altered coordination of network activity. Transcriptional and epigenetic responses converged on autism spectrum disorder (ASD) associated gene networks, with strong enrichment among high-confidence and strong-candidate ASD risk genes, suggesting that THC preferentially perturbs neurodevelopmentally vulnerable pathways. Together, these findings define a mechanistic framework in which THC disrupts early human cortical development through a cycle of transient excitatory activation, compromised structural support, and persistent epigenetic alterations, which are features specifically revealed by human PFCOs.

## Background

Cannabis is among the most commonly used psychoactive substances globally, and its use during pregnancy is increasing in parallel with global legalization and overall perceptions of safety. Prevalence studies show use of cannabis during pregnancy to be between 3% and 35% in North America^1^. This presents a significant public health concern, as Δ9-tetrahydrocannabinol (THC), the principal psychoactive cannabinoid, is lipophilic and readily crosses the placental barrier, allowing fetal exposure to cannabis use. Observational human data and preclinical evidence collectively associate prenatal cannabis exposure with adverse neonatal outcomes (i.e., low birth weight, lower grey matter volume, lower cognition) and long-term impacts on psychiatric and behavioral characteristics, such as internalizing, externalizing, attention, thought, and social problems^2^.

The mechanisms underlying these effects involve THC binding to cannabinoid receptors CB1 and CB2, which potentially disrupt associated signaling pathways that play a crucial role in fetal brain development, regulating neuronal proliferation, migration, axonal pathfinding, synaptogenesis, and circuit wiring^1^. Animal models have demonstrated that gestational THC exposure can lead to lasting alterations in gene regulation, synaptic plasticity, and epigenetic marks. For example, maternal cannabis use in rodents has been associated with long-lasting increases in histone 3 lysine 9 (H3K9) dimethylation at the dopamine D2 receptor (*Drd2*) locus, correlating with reduced *Drd2* mRNA expression in the nucleus accumbens^3^. This example suggests a potential broader epigenetic mechanism for cannabis-induced disturbances that may be relevant to addiction vulnerability and psychological disorders. In rhesus macaques, epigenome-wide studies report that prenatal THC exposure is associated with differential DNA methylation in the placenta and fetal tissues, including ∼581 CpG sites and associations with autism spectrum disorder (ASD) genes in the prefrontal cortex and other regions^4^.

Despite these collective efforts to understand the molecular basis of the effects of prenatal THC exposure, the direct transcriptomic and epigenetic consequences remain poorly understood in the human fetal brain. Several methodological challenges present barriers to such studies. Ethical and logistical considerations restrict the study of human fetal brain tissue. Additionally, animal models fail to capture human-specific aspects of cortical development, including extended neurogenic timescales, unique interneuron migratory programs, and divergent epigenetic regulation^5^. Traditional two-dimensional in vitro approaches lack the cellular diversity and three-dimensional organization required for proper neurodevelopment, with parallel comparisons showing that monolayers exhibit reduced neurogenesis efficiency and poorly defined regional identity compared to three-dimensional organoids^6^. Post-mortem tissue analyses capture only static endpoints, limited by RNA degradation, sampling constraints, and inability to track dynamic developmental trajectories^7^. Together, these constraints have precluded a mechanistic understanding of how THC disrupts human cortical development in space and time. Hence, in studying THC’s effects on the developing human brain, we have developed advanced *in vitro* models: human induced pluripotent stem cell (hiPSC)-derived brain organoids that closely mimic early human brain development. Our pre-frontal cortex Organoids (PFCOs) possess key characteristics that render them a powerful platform for neurological investigation. They recapitulate region-specific transcriptional identities, including robust expression of cortical progenitor and neuronal lineage markers such as *PAX6*, *FOXG1*, *BCL11B*, and *SATB2*, alongside supporting glial and oligodendrocytes populations marked by *GFAP*, *PDGFRA*, and *OLIG2*. PFCOs exhibit an excitatory-inhibitory neuronal balance, evidenced by co-expression of upper and deep layer markers (*CUX1*, *BCL11B*) and neurotransmitter-specific genes (*SLC32A1*, *GAD1*, *GAD2*).

We used electrophysiological and transcriptomic profiling to indicate maturation of synaptic machinery and metabolic coupling comparable to mid-gestational human cortex, allowing dynamic interrogation of early cortical network formation. Together, these attributes make PFCOs a physiologically relevant and scalable model for studying human cortical development and its perturbation by environmental exposures, such as THC. Following this, we exposed human PFCOs to THC under controlled concentration conditions to mimic prenatal exposure and profiled transcriptomic and epigenetic changes at different timepoints post-exposure. By combining bulk RNA-sequencing and whole-genome bisulfite sequencing (WGBS) data, we dissected how THC exposure remodels gene expression networks in developing cortical tissue.

## Methods

### Human induced pluripotent stem cell culture

We used **c**ontrol human induced-pluripotent stem cell (iPSC) line (Johns Hopkins University, HIRB00017883) and cultured it in T75 flasks on vitronectin-coated surfaces in Essential 8 medium (Gibco) containing DMEM/F12, L-glutamine, and sodium bicarbonate (Gibco) at 1.734 g L-1. We maintained culture in T25 flasks and passaged at a 1:2 ratio into T75 flasks every 5-7 days using Versene (Gibco) for 4 minutes at 37 °C, followed by mechanical dissociation with cell scrapers.

### Generation of prefrontal cortex organoids (PFCOs)

We generated prefrontal cortex organoids (PFCOs) from control human iPSCs using a previously published 14-day protocol that required daily media changes^8^. We washed iPSCs with 1× HBSS (Gibco) and dissociated them into single cells using Accutase^®^. We seeded cells at 10,000 cells per well in ultra-low attachment 96-well plates (300 µm, Thermoscientific) to form embryoid bodies (EBs) in Essential 8 medium supplemented with DMEM/F12, L-glutamine, sodium bicarbonate (Gibco), and the ROCK inhibitor Y-27632 (10 µM; 1:1000) for the initial 24 hours.

After 24 hours, we switched cultures to neural induction medium consisting of Neurobasal Medium supplemented with N2 and B27, containing dual SMAD inhibitors: SB431542 (10 µM; TGFβ inhibitor) and LDN193189 (0.1 µM; BMP inhibitor). We maintained EBs under these conditions for 5-7 days to promote neuroectoderm formation. We then replaced Neural induction medium with neural differentiation medium (Neurobasal Medium supplemented with N2 and B27, without SMAD inhibitors), and EBs were cultured for an additional 4-5 days to allow neural progenitor expansion.

On day 14, we embedded neural spheroids in Matrigel^®^ droplets (Product Number #356255) to provide a 3D scaffold. Subsequently, we maintained them in cerebral organoid maturation medium (Catalog #08571), which supports the emergence of diverse neural cell types and the organization of neural tissues.

We diluted THC (1.0 mg/mL in methanol (ThermoFisher); Catalog #T4764) in neurobasal media to a final concentration of 10 μg/mL^9,10^. On Day 60, we gave the PFCOs a single dose, and samples for DNA and RNA extraction were collected at 2, 6, and 24 hours after the initial THC exposure.

### RNA isolation and library preparation for bulk RNA-seq

We evaluated RNA integrity using the Agilent 4200 Tapestation, and all samples achieved RNA Integrity Number (RIN) values above 5.0. For library preparation, we used 150 ng of RNA per sample as input. We generated libraries with the NEBNext UltraExpress RNA Library Prep Kit (New England Biolabs, Ipswich, MA) according to the manufacturer’s protocol. We enriched mRNA by poly(A) selection, fragmented, reverse transcribed, and converted it into cDNA. The cDNA underwent end repair, adapter ligation, and size selection, followed by PCR amplification (11 cycles) to ensure sufficient yield while minimizing amplification bias.

We verified the library quality and size distribution on a Tapestation (Agilent) and measured concentrations using a Qubit fluorometer (Thermo Fisher Scientific). We pooled the final libraries equimolarly and sequenced on an Illumina NovaSeq X Plus using a 25B lane configuration with paired-end 150-base pair (bp, PE150) reads. Each sample achieved a minimum of 20 million reads, with an average sequencing depth of 20 million reads per sample. Base calling and demultiplexing were performed with Illumina’s DRAGEN Bio-IT platform.

### Preprocessing of sequencing data from bulk RNA-seq

We processed raw RNA-seq data using a standard pipeline that included quality control, trimming, alignment, and transcription quantification. Sequencing quality was assessed using FastQC^11^, and compiled summary reports using MultiQC^12^. We trimmed adapter sequences and low-quality bases with Trim Galore!^13^. Then we aligned the reads to a human reference genome using STAR^14^, and transcript-level quantification was performed using Salmon^15^ in alignment-based mode, generating quant.sf files for each sample. These quantifications were then imported into R using the tximport^16^ package for gene-level summarization prior to differential expression analysis.

### Bulk RNA-seq analysis in R

We identified differentially expressed genes (DEGs) using the DESeq2 package^17^ (version 1.46.0) in R^18^ (version 2025.09.0). We filtered the complete set of identified genes by a cutoff of adjusted *p*-value < 0.05 and absolute fold change > 2. Then, we subjected the DEGs to GO enrichment analysis and KEGG pathway analysis using the “clusterProfiler” R package^19^ (version 4.14.6). SFARI gene datasets were downloaded from https://gene.sfari.org/^20^.

### DNA isolation and library preparation for whole-genome bisulfite sequencing (WGBS)

Organoid samples were given to Admera Health to generate whole-genome bisulfite sequencing (WGBS) data. Genomic DNA was extracted using QIAmp DNA Micro Kit (QIAGEN, Redwood City, CA) following the manufacturer’s instructions. Isolated genomic DNA was quantified with Invitrogen™ Qubit™ Fluorometer (ThermoFisher, USA) and quality assessed with 1% Standard agarose gel (ThermoFisher, USA). Library preparation was performed using NEBNext^®^ Enzymatic Methyl-seq Kit (New England Biolabs, USA) per manufacturer’s recommendations. Briefly, gDNA was sheared to 250 bp using the Covaris S220 system (Covaris, Woburn, MA), adapters were ligated, and DNA fragments were amplified with PCR cycles. Final libraries quantity was assessed by Qubit 2.0 DNA HS Assay (ThermoFisher, USA), QuantStudio 5 System (ThermoFisher, USA), and quality was assessed by TapeStation High Sensitivity D1000 Assay (Agilent Technologies, CA, USA), QuantStudio 5 System (ThermoFisher, USA). Equimolar pooling of libraries was performed based on QC values. Samples were sequenced on an Illumina^®^ Novaseq X Plus (Illumina, California, USA) with a read length configuration of 150 PE for 600 M PE reads per sample (300M in each direction). A 20% PhiX Spike-In was added during the sequencing to ensure sequencing quality.

### Preprocessing of sequencing data from whole-genome bisulfite sequencing (WGBS)

We processed the Methyl-Seq samples (n = 9) using BiocMAP^21^, an analysis pipeline for whole-genome bisulfite sequencing (WGBS) data. Briefly, the pipeline consists of two stages – alignment and extraction. During alignment, we ran quality control (QC) checks on the raw sequencing files using FastQC^11^, followed by trimming the reads using Trim Galore!^13^, aligned to a reference genome (here, hg38) using Arioc^22^. We filtered out the low-quality or duplicate mappings. During the extraction stage, we obtained the DNA methylation proportions using *Bismark*^23^, and aggregated our results into two bsseq^24^ R objects for downstream analyses. We also produced QC metrics to examine the integrity of the processed samples. The first bsseq object contained smoothed estimates of methylated and unmethylated cytosines in CpG context across the entire genome, and the second object contained any additional cytosines in CpH context. We used only the first CpG methylated count for the present analysis.

### Differentially methylated region (DMR) Analysis

We analyzed the differentially methylated regions (DMRs) using the bsseq package in R (version 4.5.0). Prior to DMR analyses, we inspected samples for adequate coverage, defined as an average of at least 8 reads per CpG site across all samples. We also inspected various QC metrics by sample and case-control status. After dropping low-coverage CpG sites, we computed t-statistics for each case-control comparison (i.e., 2 hour versus control; 6 hour versus control) using default parameters from the *BSmooth.tstat* function in bsseq. Based on examination of the quantiles of the t-statistics, we then selected a cutoff (here, 2) to identify DMRs.

### Enrichment analysis for DMRs

Following the DMR analysis, we applied the enrichGO function from the clusterProfiler^19^ (version 4.14.6) R package to evaluate enrichment of differential methylation in Gene Ontology (GO) biological processes, molecular function, and cellular component ontologies. First, within each condition comparison, hyper- and hypo-methylated regions were isolated and tested for enrichment, separately. For each ontology, we used an FDR-adjusted p-value and q-value cut-off of 0.05 to determine significant enrichment (12 total tests). We plotted significant results as dot plots of the top 10 GO terms.

### Persistence and concordance analyses of DMRs

We also sought to examine persistent methylation changes in DMRs (using mean difference in methylation between exposure and untreated control conditions as a measure of effect size) across both group comparisons (i.e., control vs. 2 hour; control vs. 6 hour). Specifically, we assessed the overlap in genomic regions using summary output files, annotated these regions to obtain names of genes within these regions, and assessed the number of genes that overlapped across conditions. We also calculated Spearman’s rank correlations for the effect sizes (i.e., mean methylation difference) to assess stability in effects across the two timepoint conditions. We plotted the effect sizes for the top 50 overlapping genes from DMRs on a heatmap.

We also sought to examine concordance between the DMR and RNA-Seq analyses of the same samples. Similar to our assessment of persistent methylation effects, we used genomic coordinates from the summary output to identify genes within the DMRs and assessed how many of these genes also appeared as significantly differentially expressed genes (DEGs) in the RNA-Seq analysis.

### Electrophysiology

We recorded electrophysiological activity from patient-derived iPSC-based PFCOs using a MED64-Presto micro-electrode array (MEA) system. On day 65 of differentiation, we prepared the MEA plate by applying laminin and poly-D-lysine coatings, which were incubated overnight. We positioned our PFCOs on the array using a Matrigel^®^ -based hanging drop technique. We selected this approach because it leverages gravity to facilitate organoid aggregation within a controlled microenvironment, enabling spontaneous tissue-to-tissue interactions without requiring artificial scaffolding or direct physical manipulation. This technique proved particularly useful for establishing initial cellular contacts and promoting fusion under defined conditions.

For data analysis, we implemented a multi-stage preprocessing pipeline to extract high-quality spike trains from the raw recordings. Initially, we conducted bandpass filtering between 0.5 and 3000 Hz using a fourth-order Butterworth filter. We chose this filter design for its characteristically flat passband response, which preserved the integrity of physiologically relevant signals by minimizing phase distortion. The selected frequency window captured the bandwidth typically associated with neuronal action potentials and local field potential activity^8^.

## Results

### Time-resolved transcriptomic profiling shows dynamic but transient molecular responses to THC in PFCOs

We characterized the transcriptomic changes of THC treatment on the PFCOs by performing bulk RNA sequencing on the organoids harvested at 2 hours, 6 hours, and 24 hours post-THC exposure (**Figure 1A**). At each time point, we collected and sequenced three samples, for a total of 12, including three untreated control samples. We detected a total of 49,132 genes, of which 7,652, 1,886, and 360 were significantly differentially expressed (p < 0.05) at 2, 6, and 24 hours post-THC treatment, respectively. We analyzed these differentially expressed genes (DEGs) and resulting pathway enrichments to elucidate the effects of cannabinoid exposure on transcript expression in our brain models (**Figures 1C & 2**).

**Figure 1:**
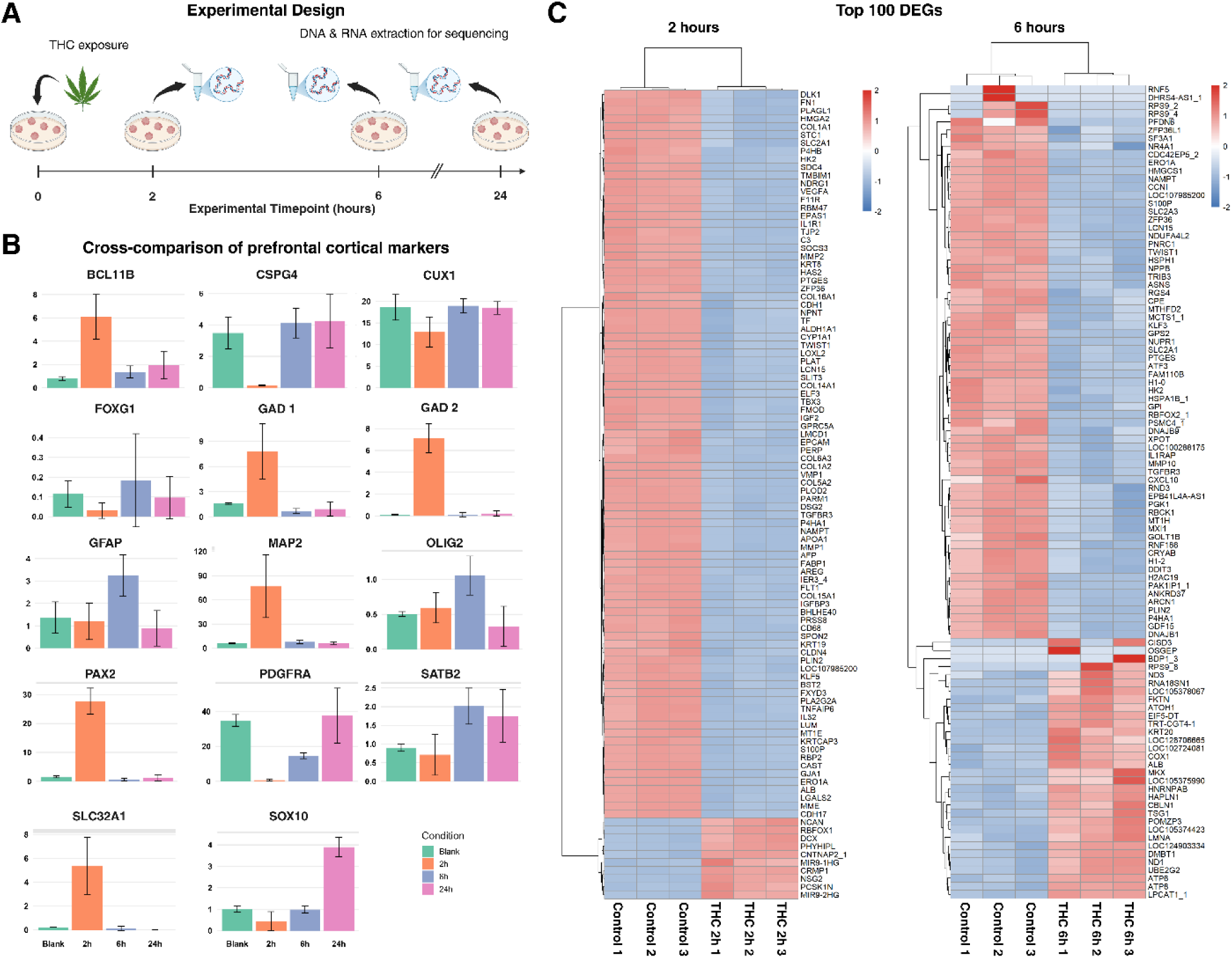
Bulk RNA-seq time-dependent transcriptomic responses to acute THC exposure in prefrontal cortex organoids. **(A)** Schematic overview of experimental design. PFCOs were exposed to THC and harvested at 2, 6, and 24 hours post-exposure. DNA and RNA were extracted for WGBS and bulk RNA-seq profiling. **(B)** Cross-comparison of canonical prefrontal cortical markers across Blank, 2h, 6h and 24h conditions. Bar heights represent transcripts per million (TPM), a normalized expression measurement that accounts for sequencing depth and gene length. **(C)** Heatmaps of the top 100 differentially expressed genes (DEGs) at 2 hours (left) and 6 hours (right), ordered by hierarchical clustering. Colors represent expression values after Z-score scaling, indicating whether a gene is expressed higher or lower relative to its own mean across samples.

To verify the regional identity of the organoids used in the THC exposure experiments, we compared expression of canonical prefrontal cortical markers across Blank, 2 hour, 6 hour, and 24 hour samples (**Figure 1B**). As expected, PFCOs robustly expressed dorsal cortical progenitor markers (*PAX6*, *FOXG1*) and lineage-defining neuronal genes (*CUX1*, *BCL11B*, *SATB2*), confirming the presence of upper- and deep-layer excitatory neuron populations. We also consistently detected inhibitory lineage markers (*GAD1*, *GAD2*, *SLC32A1*) and glial/oligodendrocyte-associated genes (*GFAP*, *PDGFRA*, *OLIG2*, *CSPG4*)^8^. Although individual markers showed time-dependent increases or decreases in transcripts per million (TPM) following THC exposure, these shifts reflected normal variability across samples rather than loss or gain of regional identity. Critically, no condition exhibited a collapse or emergence of lineage programs. With these results, we confirm that PFCOs maintain a stable prefrontal cortical transcriptional profile across all experimental timepoints, establishing the cellular context in which THC-induced transcriptional changes were subsequently measured.

### Enrichment analysis and DEGs at 2 hours are directly relevant to cannabinoid action and synaptic effects

After 2 hours of THC exposure, GO enrichment analysis revealed a strong, significant upregulation of neuronal signaling machinery, particularly synaptic organization and excitatory neurotransmission (**Figures 1 & 2**). We related upregulated biological process (BP) GO terms to development and connectivity, including terms such as forebrain development, synapse assembly, axonogenesis, regulation of neuron projection development, and regulation of synapse organization/activity. Upregulated cellular components (CC) included synaptic/postsynaptic membrane, postsynaptic specialization, neuron-to-neuron synapse, and asymmetric synapse, localizing the effects of the cannabinoid at the synapses. Functionally, we observed that the ionotropic and glutamate receptor function was upregulated. This was indicated by the molecular function (MF) GO terms such as tubule/microtubule binding, glutamate receptor activity, and ephrin receptor activity. Upregulated KEGG pathways included synaptic and neurotransmission-related pathways, such as glutamatergic synapse and axon guidance. Additionally, several pathways related to neurodegenerative diseases explicitly appear, namely Huntington’s and Parkinson’s disease. Unsurprisingly, retrograde endocannabinoid signaling pathways also demonstrated increased expression. The combination of these enrichments suggests a coordinated acute upregulation of synaptic signaling programs.

Through downregulated enrichments at 2 hours, we illustrate the suppression of structural support and adhesion pathways. Negatively enriched GO: BP terms included extracellular matrix (ECM) organization, wound healing, and regulation of adhesion. This effect was localized to the collagen-containing ECM, ER lumen, focal adhesion, and apical part of the cell as indicated by the GO:CC terms. We observed reduced expression in pathways with the GO:MF terms ECM structural constituent, cadherin and integrin binding, glycosaminoglycan binding, and growth factor binding. Downregulated KEGG pathways included ECM-receptor interaction, focal adhesion, and PI3K-Akt signaling, which align with the decreased structural functionality. This suggests a trade-off: organoids ramp up signaling but downregulate ECM integrity, possibly destabilizing their structural environment.

The specific DEGs identified in 2 hours post-THC treatment samples were strongly associated with neuronal signaling, extracellular structure, and stress response (**Figure 1C**). We observed that the growth factor-related genes, such as IGF2 and IGFBP3^25^, were among the most significantly downregulated, while *VEGFA*^26^ and *TGFBR3*^27^ also showed significant downregulation. Genes involved in ECM composition and adhesion, including *COL1A2*^28^, *COL6A3*^29^, *LUM*^30^, *SDC4*^31^, and *DSG2*^32^ were robustly suppressed, consistent with enrichment results highlighting downregulation of ECM-related pathways.

Stress- and metabolism-associated genes were also prominently differentially expressed. *NAMPT*, a key NAD^+^ biosynthetic enzyme^33^, was significantly downregulated, as was *NDRG1*, a myelination and hypoxia-responsive factor^34,35^, and *ERO1A*, which regulates oxidative protein folding in the endoplasmic reticulum^36^. Transcriptional regulators such as *BHLHE40* (hypoxia/circadian response)^37,38^ and *ZFP36*/TTP (inflammatory mRNA decay)^39^ were also downregulated. Synaptic plasticity-related genes included *PLAT* (tPA), a protease critical for NMDA receptor signaling and dendritic remodeling^40,41^.

Additional significant genes identified outside the top 100 DEGs (**Supplementary Data, DEG_2h.csv**) further support these themes. *ENHO* (adropin), a regulator of energy metabolism and cognitive function^42,43^, and *SCRT2*, a neural transcription factor required for neuronal differentiation^44^, were both upregulated. ECM remodeling was reinforced by downregulation of *VCAN* (versican)^45^ and *DSC2* (desmocollin-2)^46^, solidifying the pattern of adhesion-related suppression. Alterations in neuronal metabolism were also reflected by downregulation of *SLC2A3* (GLUT3)^47^, the primary neuronal glucose transporter, while *IQGAP1*, a cytoskeletal scaffolding protein critical for dendritic spine dynamics^48^, was reduced. Additional changes included suppression of *CD74*, a regulator of neuroinflammatory signaling^49^, and *GNRH2*, a neuroendocrine hormone linked to synaptic plasticity^50^.

Together, our data indicates that by 2 hours after THC exposure, PFCOs undergo a rapid transcriptional shift characterized by the upregulation of synaptic and neurodevelopmental programs, accompanied by the downregulation of extracellular matrix, adhesion, and metabolic support pathways. Several of these transcriptional changes overlap with pathways known to be disrupted in neurodegenerative disease and highlight potential molecular mechanisms through which THC impacts neuronal function.

### Relevant synaptic effects disappear by 6 hours, replaced by stress-related gene sets

By comparing enrichment analysis of between 2 hours and 6 hours (**Figures 1C & 2**) we observed that the initial synaptic activation effects last merely hours, and seemingly faded by 6 hours, replaced by metabolic and adaptive responses. Upregulated GO: BP included only response to xenobiotic stimulus, and for GO:CC, the terms respiratory chain complex and oxidoreductase complex. Positively enriched GO:MF gene sets included DNA helicase activity, ATP-dependent functions, and transporter/electron-transfer activity. Activated KEGG pathways were relevant to metabolism-related pathways such as oxidative phosphorylation and drug metabolism. Genes for energy production, DNA repair/replication, and detoxification were enriched, reflecting cellular adaptation to stress.

After 6 hours, THC actively suppresses protective cellular stress pathways. GO:BP terms that were downregulated included response to oxygen/hypoxia, unfolded protein response, and response to misfolded protein. The cellular locations of these effects included melanosome, focal adhesion, and ribosome, as indicated by the GO:CC terms. Affected GO:MF terms featured GTP binding/GTPase activity, nucleotide binding, and unfolded protein binding. The affected KEGG pathways included protein processing in the ER, hypoxia response, and ribosome function. These findings lead us to conclude that after the acute THC exposure, organoids downregulated key defensive mechanisms, which could potentially render them more susceptible to potential drug metabolism-related stress.

The DEGs identified at 6 hours post-THC exposure included fewer genes related to synaptic signaling than the 2 hour dataset, consistent with enrichment analyses showing a shift away from synaptic pathways (**Figure 2**). Instead, many of the observed transcriptomic changes were associated with stress responses, metabolic regulation, and structural processes. Among the genes of note were *ATP6*, which encodes a mitochondrial ATP synthase subunit, and was significantly upregulated, reflecting altered oxidative phosphorylation^51^. Several genes associated with stress and protein quality control were also present, including *PTGS2* (COX-2; **Supplementary Data, DEG_6h.csv**), an inducible enzyme linked to neuroinflammation^52^, and *CDKN1A* (p21; **Supplementary Data, DEG_6h.csv**), which regulates cell cycle arrest in response to stress^53^.

**Figure 2:**
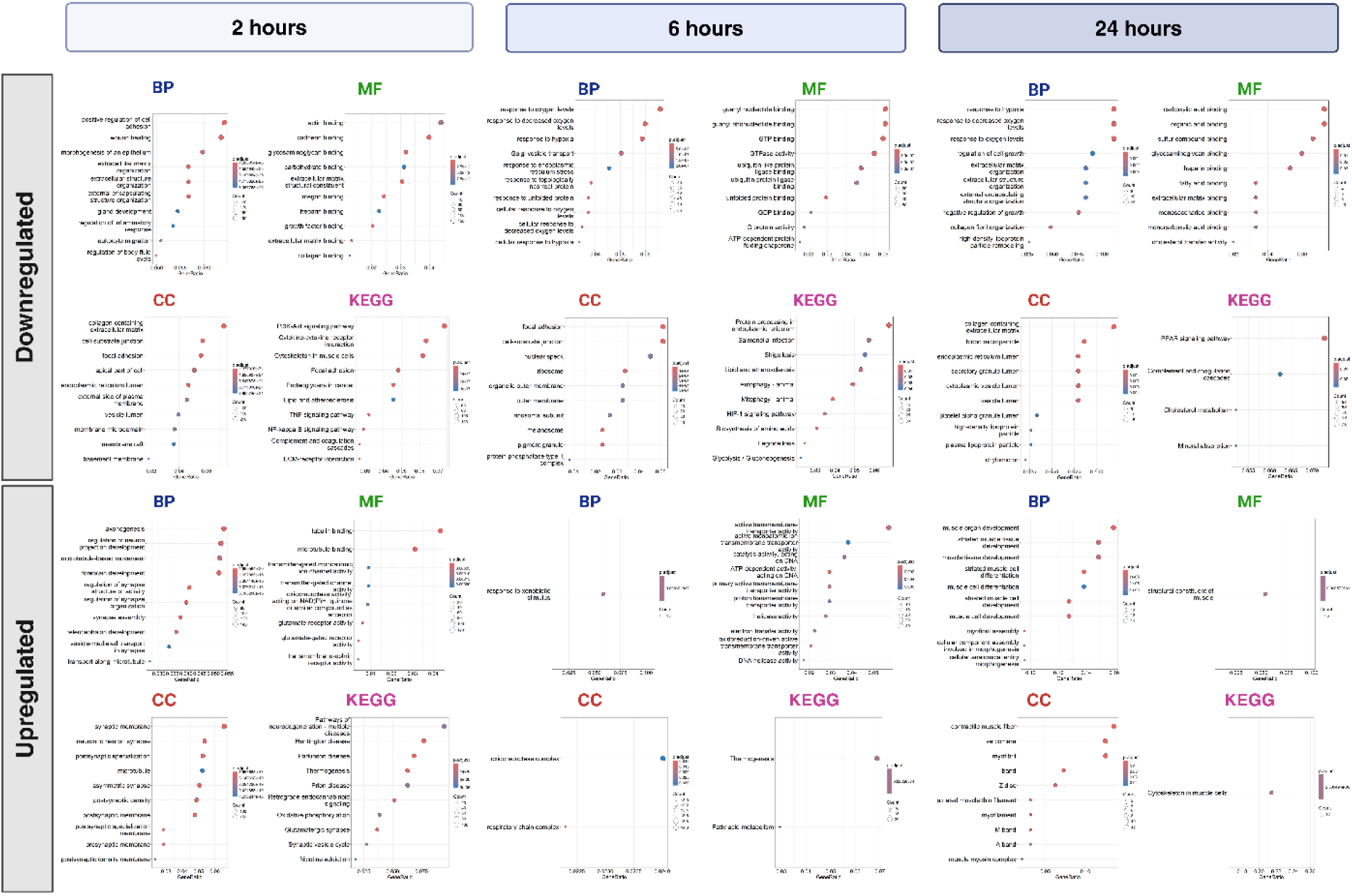
GO/KEGG enrichment of DEGs generated from bulk RNA-seq time-dependent response to THC-exposure. Results from Gene Ontology (GO) and KEGG enrichment of significantly upregulated and downregulated differentially expressed genes at 2, 6, and 24 hours post THC-exposure compared to controls. GO enrichment is separated by Biological Process (BP), Molecular Function (MF), and Cellular Component (CC) ontologies. In the dot plots, each row represents an enriched pathway, where dot color reflects the significance (adjusted p-value), dot size indicates the number of genes contributing to the enrichment, and dot position shows the enrichment ranking.

We also found that additional significant genes outside the top 100 further reinforce the dominant stress/metabolic theme. *PRPH* (peripherin), a neuronal intermediate filament protein^54^, was downregulated, suggesting possible cytoskeletal remodeling. *PHOX2A*, a transcription factor essential for autonomic nervous system development^55^, was also reduced. Genes involved in immune and inflammatory signaling were differentially expressed, such as *EREG*^56^ and *HLA-A*/*HLA-E*^57^, reflecting potential immune modulation. *NDUFA6*, a complex I mitochondrial subunit, was markedly downregulated, reinforcing the theme of impaired oxidative phosphorylation^58^.

Together, these 6 hour results highlight a transition from the strong synaptic and ECM-related transcriptional changes observed at 2 hours to a profile dominated by mitochondrial, stress, and immune-related gene alterations. This supports our hypothesis that the acute synaptic effects of THC exposure are transient. By 6 hours, the PFCOs instead show signatures of stress adaptation and metabolic imbalance, rather than direct modulation of neuronal signaling.

### Residual transcriptomic signatures at 24 hours reveal broad structural remodeling rather than sustained THC-specific effects

At 24 hours post-THC exposure, PFCOs retain only residual transcriptomic alterations, and few differentially expressed genes (DEGs) remain directly relevant to neuronal or synaptic pathways (**Figure 2**). Upregulated enrichments predominantly map to cytoskeletal and muscle-associated processes, indicating generalized structural remodeling rather than activation of true myogenic programs. These signatures span GO:BP categories such as striated muscle tissue development, myofibril assembly, and muscle organ development; GO:CC terms including sarcomere, contractile fiber, myofibril, I band, and Z disk; and GO:MF terms such as structural constituent of muscle. KEGG pathways emphasize muscle contraction and actin cytoskeleton regulation. Collectively, these findings suggest that PFCOs engage in late-phase cytoskeletal reorganization as THC-specific transcriptional activity subsides.

Downregulated enrichment profiles reinforce this interpretation. Although ECM-related pathways remain suppressed at 24 hours, the pattern broadens and becomes less THC specific, consistent with a global reduction in stress and metabolic activity. Enriched GO:BP terms include response to hypoxia, collagen fibril organization, and negative regulation of growth; GO:CC terms center on blood microparticle, ECM, and ER or secretory vesicle lumen; and GO:MF terms highlight altered organic or carboxylic acid binding, fatty acid binding, and heparin binding. Downregulated KEGG pathways, including ECM receptor interaction, lysosomal function, and metabolic processing, point to persistent impairment of structural support and intracellular homeostasis. Together, these enrichments indicate that PFCOs enter a broader low-activity state marked by continued ECM and metabolic suppression rather than sustained THC-responsive signaling.

Individual DEGs from the **Supplementary Data (DEG_24h.csv)** show a similar pattern. Only a small subset exhibits neurodevelopmental relevance at this later time point. *NDN* (Necdin), which supports neuronal survival and differentiation and is implicated in Prader–Willi syndrome^59,60^, is downregulated. Several metallothionein genes (*MT1H*, *MT1E*, *MT1G*), key regulators of oxidative stress responses relevant to Alzheimer’s and Parkinson’s disease^61^, are also suppressed. Downregulation of *P4HA1*, required for collagen hydroxylation^62^, signals persistent ECM dysregulation. *NAV3*, involved in axon guidance and neuronal migration^63^, is upregulated, while *GLRA2* (glycine receptor alpha 2 subunit) shows increased expression, pointing to subtle effects on inhibitory neurotransmission^64^. *HK2*, a regulator of glycolytic flux and energy metabolism in brain cells^65^, is reduced, suggesting emerging metabolic vulnerability. Immune-related alterations persist as well, including suppression of *C1QB*, a complement component implicated in synaptic pruning and neuroinflammation^66^. Changes in cytoskeletal genes such as *ACTC1*^67^ and *TTN*^68^ likely reflect broad structural remodeling rather than neuron-specific programs.

Overall, the 24 hour transcriptomic profile demonstrates that THC-induced molecular responses attenuate substantially with time. Residual effects primarily encompass stress-response attenuation, metabolic suppression, ECM disruption, and cytoskeletal reorganization. Only a minimal set of DEGs retains ties to neuronal biology, indicating that PFCOs largely resolve early THC specific transcriptional programs by this later stage. At 2 hours, enrichment analysis revealed strong THC relevant signatures, with upregulated pathways concentrated in synaptic and neuronal processes and downregulated pathways dominated by extracellular matrix and adhesion. By 6 hours, the profile shifted toward metabolic and stress-response pathways, indicating an early transition away from direct neuronal signaling effects. At 24 hours, enrichment signals weakened further and were clearly less THC specific. Upregulated terms centered on muscle and cytoskeletal remodeling, while downregulated terms continued to reflect broad suppression of ECM and stress-related pathways. Notably, ECM and adhesion pathways were consistently downregulated at both 2 hours and 24 hours, suggesting a persistent reduction in extracellular support structures across the exposure window. We interpret the reduced pathway specificity at 6 hours and especially at 24 hours as evidence of secondary or compensatory responses, consistent with a transient primary effect of THC exposure followed by broader remodeling and recovery processes in PFCOs.

### Whole genome methylation sequencing reveals differentially methylated regions in response to THC exposure, with methylation in genes implicated in autism spectrum disorder (ASD)

We also sought to characterize the epigenetic changes of THC treatment on the PFCOs, performing whole-genome bisulfite sequencing (WGBS) on the organoids harvested at 2 hours, 6 hours post-THC exposure, and untreated controls. We detected 29,401,795 CpG sites across all chromosomes (including X, Y, and mtDNA). Among these, 8,811,546 CpG sites had an average coverage of at least 8 reads across all samples and were included in the differentially methylated region (DMR) analysis. Mean coverage across cases (M = 11.9) and controls (M = 13.4) were similar, *t (*6.9) = 1.57, *p* = 0.15. All samples had an error rate for primary mappings <= 1%; similar mean fragment length (cases: M = 139.8; controls: M = 137.9), *t* (4.3) = −0.38, *p* = 0.71; and similar number of concordant pairs (cases: M = 304,034,353; controls: M = 312,732,167), *t* (3.6) = 0.46, *p* = 0.66. The mean percent methylated CpG sites were also similar between cases (M = 68.8%) and controls (M = 67.36), *t*(5.2) = −1.36, *p* = 0.22. (**Figure 3 & 4A-D**). ***DMR analysis***

**Figure 3:**
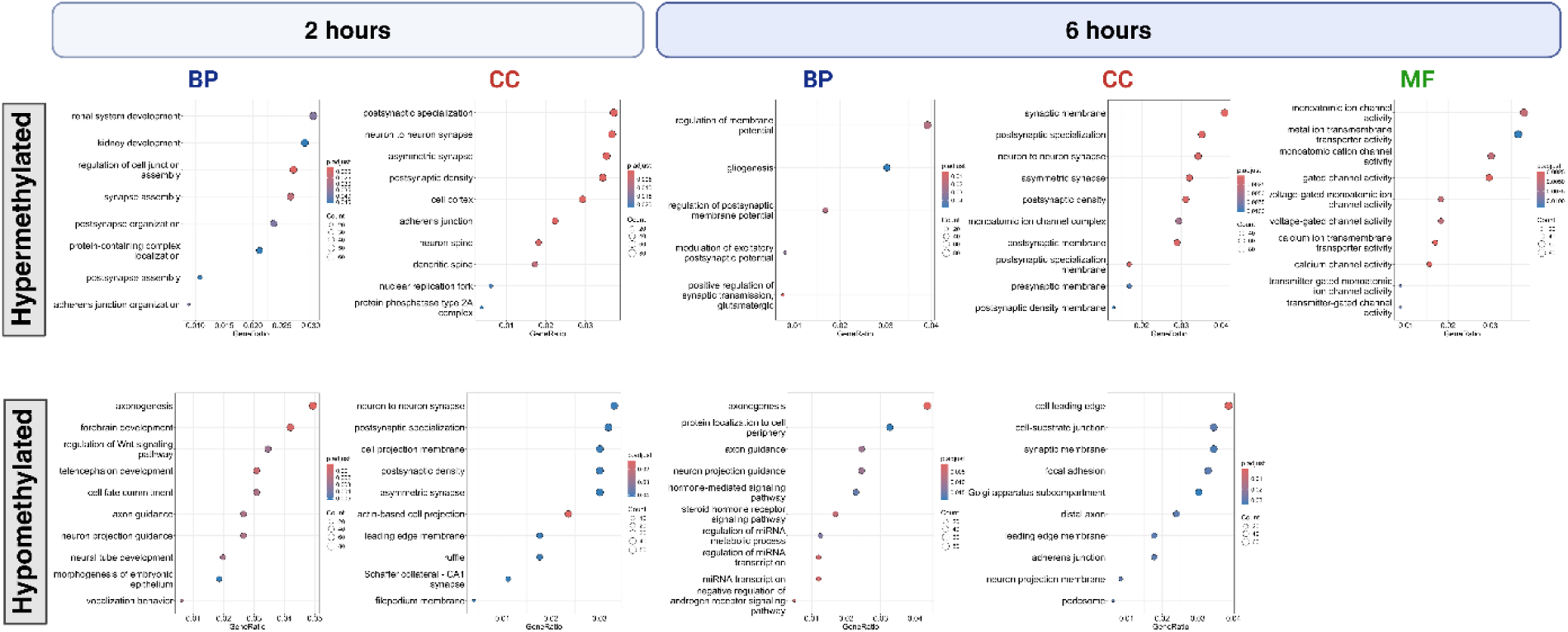
GO enrichment generated from WGBS time-dependent response to THC-exposure. Results from WGBS DMR GO enrichment analyses for Biological Process (BP), Molecular Function (MF), and Cellular Component (CC) ontologies for 2 and 6 hour conditions, compared to controls, split by significant hyper- and hypo-methylated regions. In the dot plots, each row represents an enriched pathway, where dot color reflects the significance (adjusted p-value), dot size indicates the number of genes contributing to the enrichment, and dot position shows the enrichment ranking.

Differential methylation analysis identified genomic regions where cytosine methylation levels changed in response to THC exposure, providing an insight into early epigenomic mechanisms that may have preceded or shaped transcriptional responses. Using an absolute t-statistic cutoff of 2, we detected 7,107 DMRs at 2 hours relative to the untreated control condition. At 2 hours post-THC exposure, we observed clear evidence of such regulation. The majority of DMRs, 52%, were hypermethylated, while the remaining 48% were hypomethylated compared to controls. This balanced distribution indicated that THC alters both gene-suppressive and gene-permissive methylation states.

We illustrate the genomic distribution of these DMRs in **Figure 4A-D**. Most DMRs, approximately 41%, occurred within promoter regions, underscoring their potential to directly influence transcriptional initiation. A substantial fraction also mapped to intronic regions (36%) and distal intergenic elements (16%), suggesting that THC-induced methylation changes may have extended to enhancers and other long-range regulatory sites. Together, these patterns demonstrated that early THC exposure produced rapid, locus-specific shifts in the PFCO epigenome that modified multiple layers of gene regulatory architecture.

**Figure 4:**
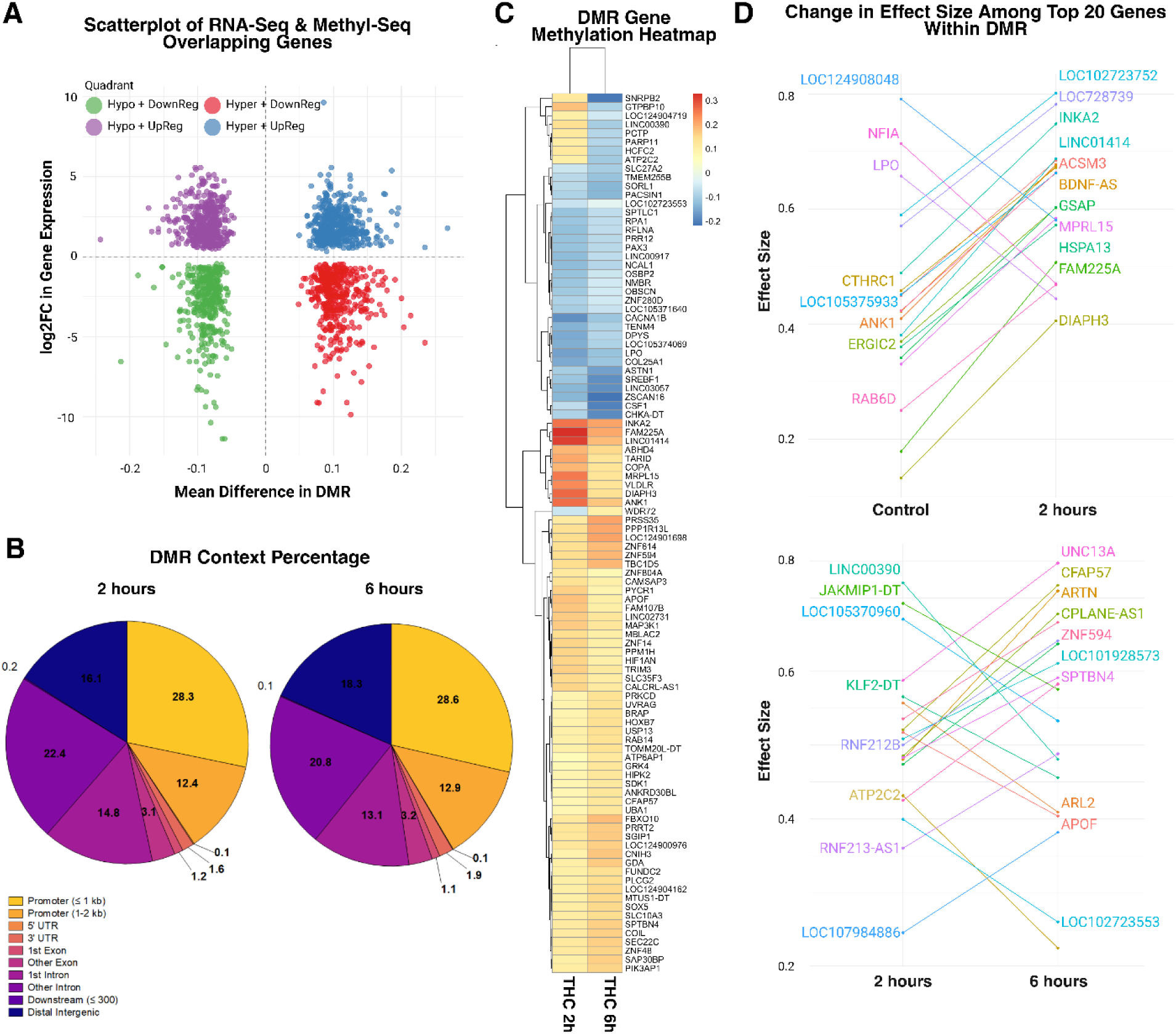
Graphical representation of significant differentially methylated regions, overlap with RNA-Seq results, and gene-level result. **(A)** Overlap in effect size and direction between overlapping genes in methyl-seq and RNA-seq analyses. **(B)** Description of gene context among differentially methylated regions (DMRs) in 2 hour and 6 hour conditions, respectively. **(C)** Heatmap of differential methylation, compared to controls, among top 100 genes in significantly differentially methylated regions overlapping across the 2 and 6 hour conditions. **(D)** Top panel: Largest effect size changes among the top 20 genes mapped to significantly differentially methylated regions. Bottom panel: Largest effect size changes among the top 20 genes mapped to significantly differentially methylated regions that overlapped between 2 and 6 hour conditions.

When comparing the control samples to those at 6 hours, we detected 7,560 DMRs, which were slightly more than at 2 hours. Similarly, the majority (52%) of DMRs were hypermethylated at 6 hours compared to untreated controls. Similar to the 2 hour post-THC exposure, the majority of DMRs (∼41%) were located within promoter regions, with DMRs also in intronic (34%) and distal intergenic regions (∼18%). We classified each DMR according to its genomic feature, defined as annotated elements such as promoters, introns, exons, UTRs, and distal intergenic regions. This allowed us to assess whether THC-induced methylation changes concentrated in regulatory regions or were distributed more broadly across the genome (**Figure 4)**.

### Enrichment analyses

We plotted the GO enrichment analyses of significant DMRs at 2 hours and 6 hours, separately, split by hyper- and hypomethylated regions (**Figure 3**). Two hours after exposure, we found significant enrichment of hypermethylation in pathways related to synapse formation and assembly, as well as postsynaptic structure. This suggested an epigenetic suppression and potential reduced transcriptional potential of genes involved in synaptogenesis and cell adhesion in the developing brain. We also detected enrichment of hypomethylation in pathways related to neurodevelopment. Notably, there was evidence of enrichment in the Wnt signaling pathway, which plays key roles in cell fate determination and differentiation during prenatal development^69^. Together, these findings lead us to suggest that THC exposure creates a paradoxical state, where hypomethylation in some regions promotes early-stage neuronal growth and differentiation while hypermethylation in other regions simultaneously inhibits the maturation and stabilization of synapses.

Enrichment results after 6 hours were similar, with enrichment of hypomethylation in pathways related to axonogenesis and axon and neural projection guidance, suggesting a sustained push for neuronal growth and connection. Interestingly, enrichment of hypermethylation at 6 hours shifted from pathways related to synapse assembly to pathways involved in electrical function of synapses. Specifically, enrichment in pathways related to membrane potential and excitatory postsynaptic potential suggested suppression of neuronal firing and communication. Combined with the 2 hour enrichment results, we infer that although THC exposure may contribute to increased neuronal growth and development, over time these circuits become impaired and functionally silent.

### Persistence of effects

We next sought to examine the persistence of effects in two cases: control to 2 hours and from 2 hours to 6 hours post-THC exposure. That is, we wanted to know whether methylation changes observed at 2 hours persisted at 6 hours or returned to untreated levels as in the control. We plotted the top 20 genes within DMRs that showed the greatest change in effect size (i.e., mean methylation difference) between the control and 2 hour conditions. (**Figure 4D**) We observed that many top genes were located within regions with hypermethylated CpG sites compared to untreated controls. Interestingly, *NFIA* was among the top genes in regions marked by hypomethylation at 2 hours. *NFIA*, located on chromosome 1, is a protein-coding member of the nuclear factor 1 family of transcription factors^70^. Because several THC-responsive genes have neurodevelopmental relevance, we cross-referenced these findings with the Simons Foundation Autism Research Initiative (SFARI) gene database^20^, a curated resource of genes associated with autism spectrum disorder. *NFIA* is classified by SFARI as a strong ASD-associated candidate (Category 2). We then selected genes within significant DMRs that overlapped between the 2 hour and 6 hour conditions to determine how methylation levels changed or may have returned to untreated levels. As shown in **Figure 4**, the top 20 genes at these overlapping time points did not align with the top genes from the 2 hour versus control comparison. Similar to the earlier contrast, the majority of the strongest effect of size changes reflected hypermethylation from 2 hours to 6 hours. One gene in this overlapping set, *UNC13A*, was consistently hypermethylated across conditions. When evaluated in the context of our prior SFARI comparison, UNC13A also appears in the database as a strong ASD-associated candidate.

### Electrophysiological characterization of PFCO following THC exposure

We analyzed four key parameters of neuronal activity in PFCOs exposed to THC compared to untreated controls (Control) across three timepoints: 2, 24, and 48 hours post-exposure. We plotted the electrophysiological activity of PFCOs following acute THC exposure (**Figure 5A**). At 2 hours post-exposure, we observed that the THC group exhibited a significant reduction in spike frequency relative to the control, accompanied by a marked increase in inter-spike interval (ISI), indicating transiently reduced firing activity. By 24 and 48 hours, both metrics recovered, and the spike frequency and ISI were comparable between groups. Burst rate remained similar across conditions at all timepoints. In contrast, we observed that the burst duration showed greater variability: the THC group displayed a tendency toward longer burst durations at 24 hours, though this effect was not sustained and diminished by 48 hours. Hence, we concluded that THC induced an early but transient suppression of network firing, while longer-timescale burst properties are more variable and less consistently affected.

**Figure 5:**
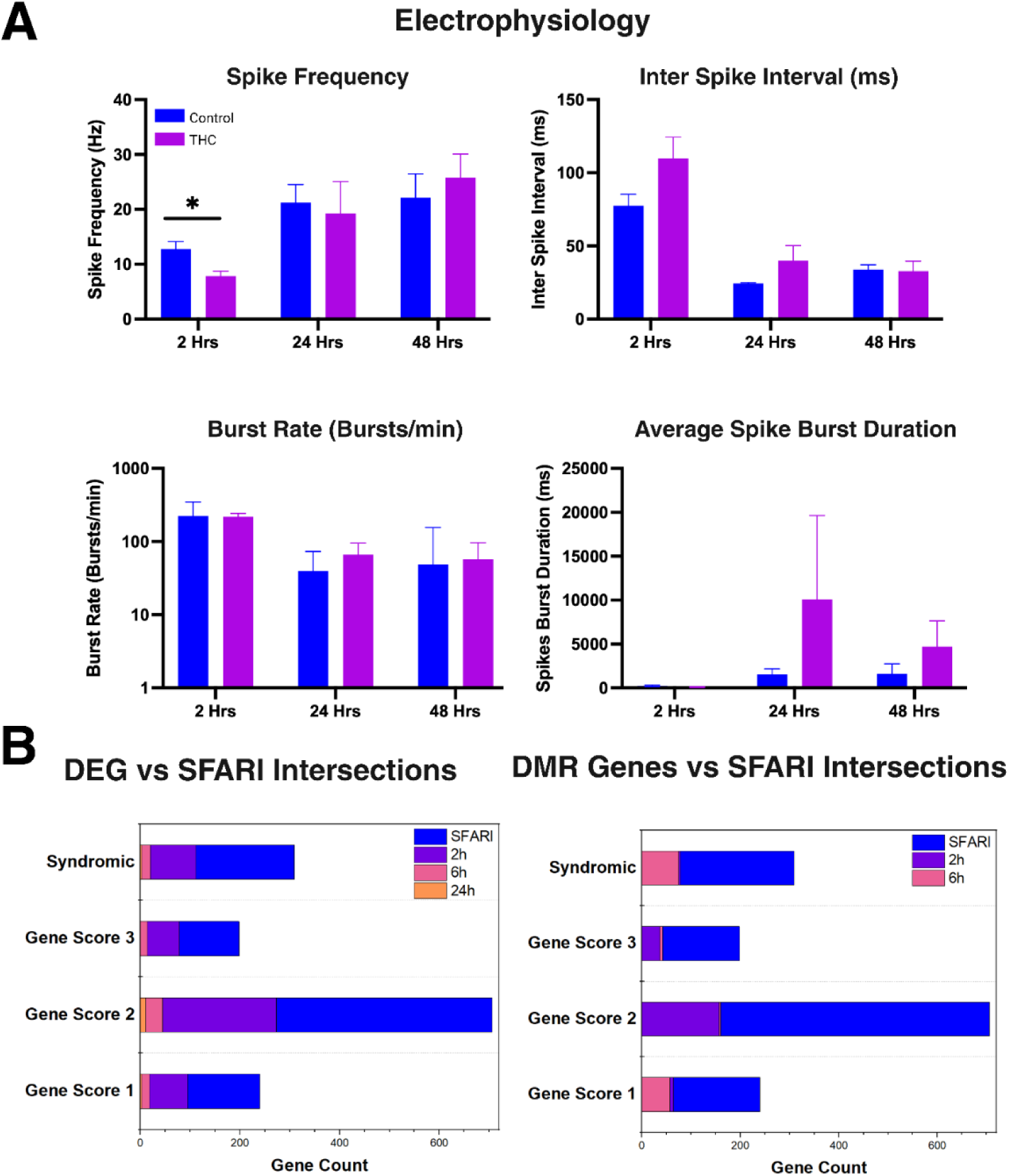
Electrophysiological measurements and SFARI gene intersection. **(A)** Electrophysiological activity of PFCOs exposed to THC or vehicle controls (Control) at 2, 24, and 48 hours. Spike frequency, ISI, burst rate, and burst duration were compared using unpaired two-tailed t-tests. A significant increase in spike frequency was observed at 2 hours (p < 0.05); no other measures showed significant differences. Data are mean ± SEM. **(B)** Intersection of THC-responsive genes with autism spectrum disorder (ASD)-associated genes from the SFARI Gene database. Bars are overlapping and the rightmost point of each color represents the value of gene count for that category. Left: Overlap between differentially expressed genes (DEGs) at 2, 6, and 24 hours and SFARI categories. Right: Overlap between differentially methylated region (DMR)-associated genes at 2 and 6 hours and the same SFARI categories. SFARI categories are the following: Gene Score 1: High Confidence, Gene Score 2: Strong Candidate, Gene Score 3: Suggestive Evidence, and Syndromic.

### THC alters gene networks enriched for high-confidence autism risk genes

To assess whether THC exposure perturbs molecular pathways relevant to neurodevelopmental disorders, we examined its relationship to autism spectrum disorder (ASD)–associated genes. ASD was selected because many early neurodevelopmental pathways disrupted in our data overlap with pathways implicated in ASD risk. We therefore intersected our RNA-seq DEGs and WGBS DMR-associated genes with curated ASD-risk genes from the SFARI Gene database (**Figure 5B**). SFARI Gene^20^ compiles ASD-associated genes from human genetic studies, and each gene was assigned a Gene Score based on the strength of evidence: **Gene Score 1 (High Confidence):** Genes clearly implicated in ASD, typically supported by ≥3 de novo likely gene-disrupting mutations (LGDs). Many met genome-wide significance or an FDR < 0.1 in large sequencing studies. **Gene Score 2 (Strong Candidate):** Genes with two reported de novo LGDs, or genes uniquely implicated by replicated genome-wide associated studies where the associated risk variant showed functional impact. **Gene Score 3 (Suggestive Evidence):** Genes with one known de novo LGD, or those supported by emerging association or inheritance evidence that has not yet met a rigorous statistical threshold. **Syndromic (S):** Genes where mutations caused a syndrome for which ASD frequently co-occurs. Some of these may also carry independent evidence for idiopathic ASD (not mutually exclusive with Gene Scores 1-3). Across the SFARI database, this corresponds to 240 High Confidence (Score 1), 706 Strong Candidate (Score 2), 198 Suggestive (Score 3), and 309 Syndromic genes.

### THC-responsive DEGs show strong early overlap with high-confidence ASD-risk genes

At just 2 hours post-exposure, we observed many High Confidence (Category 1, **Figure 5B**) genes: 95 genes, nearly 40% of the entire SFARI High Confidence set, altered at the transcript level. We saw the strongest enrichment among Category 2 Strong Candidate genes, with 273 DEGs overlapping this group. We also saw a considerable overlap in the syndromic genes (111 genes), indicating that THC perturbs transcriptional programs associated not only with idiopathic ASD but also with broader neurodevelopmental syndromes.

By 6 hours, the number of intersecting DEGs declined but remained notable, and by 24 hours, only a handful of SFARI genes remained differentially expressed. This time-dependent contraction suggested that THC induces a sharp but transient early wave of transcriptional dysregulation that disproportionately affects ASD-relevant genes.

### THC-induced DMR-associated genes converge on the same ASD-risk categories

We saw a similar trajectory in epigenetic intersections but with greater temporal stability. THC-induced DMR-associated genes at 2 hours overlapped extensively with SFARI categories, including 64 High Confidence and 157 Strong Candidate genes. Importantly, unlike the transcriptional changes that diminished by 24 hours, the epigenetic overlap remained highly consistent between 2 and 6 hours, with Strong Candidate overlaps remaining nearly identical (157 vs. 160 genes). This indicated that while gene expression changes represent a rapid but transient response, early methylation changes occur quickly and remain stable during the initial exposure window.

A consistent pattern emerged across datasets: THC exposure disrupted both transcriptional and epigenetic regulation of genes with well-established or strongly suggested genetic links to ASD. The particularly strong enrichment of Category 2 Strong Candidate genes highlighted a coordinated impact on pathways supported by robust but still-evolving genetic evidence. Together, these findings revealed that ASD-associated gene networks represented early molecular targets of THC in developing PFCOs with immediate transcriptional responses followed by more sustained epigenetic alterations.

## Discussion

This study provides a multi-layered view of how acute THC exposure disrupts early human cortical development, integrating transcriptomic, epigenetic, and electrophysiological results from hiPSC-derived prefrontal cortex organoids (PFCOs). Across modalities, a cohesive mechanistic picture emerged: THC induced an immediate excitatory shift in immature cortical circuits, weakened extracellular and structural support systems, and imposed epigenetic constraints on synaptic maturation. These findings offer an insight into how early cannabinoid exposure may alter developmental trajectories during a critical window of human cortical formation.

### Early synaptic activation without structural or epigenetic support

Within 2 hours post-THC exposure, our PFCOs displayed strong upregulation of genes involved in synaptic organization, glutamatergic signaling, axonogenesis, and cortical development. This was consistent with established roles of CB1 receptors in enhancing presynaptic glutamate release, modulating neurotransmitter vesicle cycling, and influencing early synaptic plasticity^71,72^. Similar acute activation of excitatory pathways have been observed in rodent prenatal cannabis exposure studies, where THC transiently increases neuronal excitability and modifies synaptic signaling^73^.

However, this excitatory upregulation occurred simultaneously with robust suppression of ECM organization, adhesion, and collagen-related pathways, suggesting a reduced capacity for stabilizing newly activated circuits. The ECM is recognized as a critical regulator of neuronal migration, synaptogenesis, and plasticity^74^. This suggests that disrupting ECM components during development can destabilize excitatory-inhibitory balance and impair dendritic spine maturation. Thus, THC appears to produce a mismatch: neurons are pushed towards early activation without receiving the structural scaffolding necessary to consolidate synapses.

### Epigenetic remodeling reinforces delayed destabilization

The WGBS data further supported this mismatch model. At the same early timepoint (2 hours), hypermethylation targeted promoters of genes involved in synapse assembly and postsynaptic specialization, indicating epigenetic repression of synaptic stabilization. At the same time, hypomethylation enriched for Wnt signaling and neurodevelopmental pathways suggests permissiveness towards neuronal differentiation. Wnt signaling plays critical roles in cortical progenitor proliferation and neuronal fate specification^75^. Similar dual effects promoting differentiation while weakening maturation have been described in primate placenta and fetal cortex following prenatal cannabis exposure. In rhesus macaques, THC exposure induced >500 differentially methylated CpGs in fetal brain, many mapping to neurodevelopmental and ASD-linked genes^4^. Together, these data suggest that THC activates networks before they are structurally ready and recruits epigenetic marks that limit maturation, potentially creating an unstable developmental trajectory.

### Temporal cascade: from early excitatory bias to metabolic and stress response

By 6 hours, transcriptomic signatures shifted sharply from synaptic processes to stress- and metabolism-related pathways, including oxidative phosphorylation, DNA repair, and xenobiotic metabolism. Cannabinoids have been previously shown to modulate mitochondrial activity, disrupt calcium homeostasis, and impair oxidative metabolism^76^. Upregulation of detoxification pathways and suppression of unfolded protein and hypoxia responses suggest that early THC-induced signaling places metabolic burden on developing neurons.

Epigenetically, hypomethylation was enriched in axonogenesis and guidance pathways, demonstrating a push for neuronal growth, while hypermethylation targeted genes involved in synaptic electrical function, such as excitatory postsynaptic potential. These findings suggest that although neurons may continue to grow and form new connections, their functional excitability is limited by epigenetic regulation. In animal models, THC exposure has been associated with persistent impairments in long-term potentiation and altered excitability in cortical and hippocampal circuits^77^.

### Residual and disease-relevant effects at 24 hours

At 24 hours, transcriptional effects were diminished but included downregulated of NDN (Necdin), a gene critical for neuronal survival and implicated in Prader-Willi syndrome^59,60^, as well as metallothionein genes important for oxidative stress defense. Although differential expression was dominated by MT-1 isoforms, it is worth noting that MT-3 has previously been implicated in neuromodulatory events and pathogenesis of Alzheimer’s disease^61^. These changes suggest lingering impairments in neuronal resilience and stress handling, with possible relevance to long-term neurodevelopmental risk. While WGBS was not extended to the 24 hour condition in this study, the persistence of ECM suppression and immune-related changes at the transcriptomic level implies that epigenetic mechanisms may underlie the durability of these effects.

### Delayed electrophysiological disruptions reflect earlier molecular instability

Electrophysiology provides functional validation of this temporal model. 24 hours post THC-exposure, PFCOs displayed dramatically prolonged burst durations without changes in spike frequency or burst rate. This phenotype suggests disruptions in burst termination, synchronization, or inhibitory balance rather than simple hyperexcitability. Burst prolongation has been observed in models of disrupted ECM, impaired GABAergic refinement, and early synaptic destabilization, aligning with the molecular findings of ECM suppression, altered inhibitory genes, and synaptic methylation signatures^78^. We observed that by 48 hours, burst metrics normalized. However, the persistant behavior of ECM suppression and stable DMRs at synaptic and ASD-associated loci suggests that longer-term of repeated exposures could produce cumulative network alterations that would not resolve as quickly.

### Engagement of ASD-related gene networks highlights a vulnerable developmental window

One of the most compelling findings is the early enrichment of THC-responsive genes among curated ASD-risk genes in the SFARI database^20^. We observed that nearly 40% of High Confidence ASD genes were differentially expressed at 2 hours, along with hundreds of Strong Candidate and Syndromic genes. While transcriptional overlaps diminished by 6-24 hours, epigenetic intersections remained remarkably stable. Several notable genes, including *NFIA* and *UNC13A*, carried strong ASD-related evidence and played established roles in cortical differentiation and synaptic release^79,80^. These results parallel findings in cannabis-exposed primates and human placenta, where DMRs are enriched near ASD-implicated genes^4^. Although this study does not claim causation, the agreement across species suggests that ASD-relevant gene networks are disproportionately sensitive to prenatal cannabinoid exposure.

### Strengths, limitations, and directions for future research

This work leverages a region-specific human organoid model, multi-omic integration, and electrophysiology recordings to dissect THC effects across several biological layers. However, important limitations remain. Human exposure is typically chronic and involves fluctuating THC levels. Chronic exposure studies may reveal cumulative or non-linear effects, especially at the epigenetic level. Additionally, bulk RNA-seq and WGBS mask cell-type-specific effects, particularly in cortical progenitors, excitatory neurons, interneurons, and glia. Single-cell bisulfite sequencing, scATAC-seq, or CITE-seq approaches would greatly enhance mechanistic insight. Furthermore, PFCOs recapitulate mid-gestational cortex but cannot model placental transfer, maternal immune environment, hormonal context, or long-range cortical connectivity. Further studies should incorporate chronic THC modelling, multi-week recordings, and microglia- or vascular-enhanced models to better recapitulate *in vivo* neurodevelopment. Direct gene editing of key THC-responsive loci (such as *NFIA*, *UNC13A* or *NDN*) may help determine causal contributions. Integration with single-cell methylation and spatial transcriptomics would also clarify which populations are most vulnerable.

## Conclusions

Together, these findings support a mechanistic framework in which THC induces a rapid excitatory push in the developing cortex, suppresses extracellular and adhesion structures needed for stable synaptogenesis, and recruits epigenetic marks that limit synaptic maturation. This imbalance produces a delayed disruption in the timing and duration of coordinated neuronal firing events and engages gene networks with strong relevance to ASD and neurodevelopmental disease. These results highlight the molecular and functional vulnerabilities of developing neuronal networks to cannabinoid exposure and underscore the importance of understanding prenatal cannabis use in the context of early brain development.

## List of abbreviations

THC: Δ9-tetrahydrocannabinol
iPSC: induced pluripotent stem cell
PFCO: prefrontal cortex organoid
RNA-seq: RNA sequencing
WGBS: whole-genome bisulfite sequencing
ASD: autism spectrum disorder
CB1: cannabinoid receptor 1
CB2: cannabinoid receptor 2
H3K9: histone H3 lysine 9
Drd2: dopamine D2 receptor
mRNA: messenger
RNA EBs: embryoid bodies
HBSS: Hank’s Balanced Salt Solution
ROCK: Rho-associated coiled-coil kinase
SMAD: Sma/MAD family of signaling proteins
BMP: bone morphogenetic protein
DMEM/F12: Dulbecco’s Modified Eagle Medium / Ham’s F-12
RIN: RNA Integrity Number
cDNA: complementary
DNA QC: quality control
PE150: paired-end 150 bp sequencing
DRAGEN: Dynamic Read Analysis for GENomics
STAR: Spliced Transcripts Alignment to a Reference
DEGs: differentially expressed genes
GO: Gene Ontology
KEGG: Kyoto Encyclopedia of Genes and Genomes
SFARI: Simons Foundation Autism Research Initiative
gDNA: genomic DNA
PCR: polymerase chain reaction
PE: paired-end
hg38: Human Genome version 38
CpG: cytosine-phosphate-guanine
CpH: cytosine followed by a non-guanine base
DMR: differentially methylated region
UTR: untranslated region
FDR: false discovery rate
MEA: micro-electrode array
Hz: hertz
TPM: transcripts per million
BP: Biological Process
CC: Cellular Component
MF: Molecular Function
tPA: tissue plasminogen activator
NMDA: N-methyl-D-aspartate
ATP: adenosine triphosphate
GTP: guanosine triphosphate
COX-2: cyclooxygenase-2
HLA: human leukocyte antigen
mtDNA: mitochondrial DNA
Wnt: Wingless-related integration site
ECM: extracellular matrix
NAD+: nicotinamide adenine dinucleotide
GLUT3: glucose transporter 3
tRNA: transfer RNA

## DECLARATIONS

### Ethics approval and consent to participate

Human induced-pluripotent stem cell line usage was reviewed and approved by the Johns Hopkins Institutional Review Board under protocol HIRB00017883. All lines were originally derived with informed donor consent and institutional ethical approval at the source institutions.

### Consent for publication

Not applicable.

### Availability of data and materials

All data supporting the findings of this study are available within the paper and its Supplementary Information. GEO accession numbers for the RNA-seq and WGBS datasets are pending approval and will be provided upon release.

## Author contributions

L.D.O., D.S., A.K., and B.M. conceptualized the study. K.J., A.P., and A.K. performed experiments to collect data. L.D.O., D.S., K.J., A.K., and B.M. conducted formal analysis, data curation, and interpretation on acquired data. L.D.O., D.S., A.P., and A.K. wrote the manuscript. A.K. and B.M. supervised the project. All authors read and approved the final manuscript.

## Supporting information

2 hours Post-THC Treatment DEGs

6 hours Post-THC Treatment DEGs

24 hours Post-THC Treatment DEGs

2 hours Post-THC Treatment DMR Genes

6 hours Post-THC Treatment DMR Genes

## Acknowledgements

All figures were assembled and created with BioRender.com. Kathuria, A. (2026) https://BioRender.com/b1wlq0d. Organoid samples were provided to Admera Health for WGBS generation.

## Supplementary Materials

- DEG_2h.csv
- `o` **Title:** 2 hours Post-THC Treatment DEGs
- `o` **Description:** List of significant DEGs 2 hours post-THC treatment. Includes average normalized expression (baseMean), estimated log2 fold change (log2foldChange), standard error of the estimated log2 fold change (lfcSE), Wald statistic (stat), unadjusted p-value (pvalue), and p-value adjusted for multiple testing (padj).
- DEG_6h.csv
- `o` **Title:** 6 hours Post-THC Treatment DEGs
- `o` **Description:** List of significant DEGs 6 hours post-THC treatment. Includes average normalized expression (baseMean), estimated log2 fold change (log2foldChange), standard error of the estimated log2 fold change (lfcSE), Wald statistic (stat), unadjusted p-value (pvalue), and p-value adjusted for multiple testing (padj).
- DEG_24h.csv
- `o` **Title:** 24 hours Post-THC Treatment DEGs
- `o` **Description:** List of significant DEGs 24 hours post-THC treatment. Includes average normalized expression (baseMean), estimated log2 fold change (log2foldChange), standard error of the estimated log2 fold change (lfcSE), Wald statistic (stat), unadjusted p-value (pvalue), and p-value adjusted for multiple testing (padj).
- DMR_2h_genes.csv
- `o` **Title:** 2 hours Post-THC Treatment DMR Genes
- `o` **Description:** List of significant DMR-associated genes 2 hours post-THC treatment. Includes average methylation level of the control PFCOs (control), average methylation level for the PFCOs 2 hours post THC-treatment (twoHour), and mean difference in methylation between the two groups (meanDiff).
- DMR_6h_genes.csv
- `o` **Title:** 6 hours Post-THC Treatment DMR Genes
- `o` **Description:** List of significant DMR-associated genes 6 hours post-THC treatment. Includes average methylation level of the control PFCOs (control), average methylation level for the PFCOs 6 hours post THC-treatment (sixHour), and mean difference in methylation between the two groups (meanDiff).

## References

1. Nashed MG, Hardy DB, Laviolette SR. Prenatal Cannabinoid Exposure: Emerging Evidence of Physiological and Neuropsychiatric Abnormalities. Front Psychiatry [Internet]. 2021 Jan 14 [cited 2025 Sept 25];11. Available from: https://www.frontiersin.org/journals/psychiatry/articles/10.3389/fpsyt.2020.624275/full

2. Paul SE, Hatoum AS, Fine JD, Johnson EC, Hansen I, Karcher NR, et al. Associations Between Prenatal Cannabis Exposure and Childhood Outcomes: Results From the ABCD Study. JAMA Psychiatry. 2021 Jan 1;78(1):64–76.

3. Morris CV, DiNieri JA, Szutorisz H, Hurd YL. Molecular mechanisms of maternal cannabis and cigarette use on human neurodevelopment. Eur J Neurosci. 2011 Nov;34(10):1574–83.

4. Shorey-Kendrick LE, Roberts VHJ, D’Mello RJ, Sullivan EL, Murphy SK, Mccarty OJT, et al. Prenatal delta-9-tetrahydrocannabinol exposure is associated with changes in rhesus macaque DNA methylation enriched for autism genes. Clin Epigenetics. 2023 July 6;15(1):104.

5. Zhao X, Bhattacharyya A. Human Models Are Needed for Studying Human Neurodevelopmental Disorders. Am J Hum Genet. 2018 Dec 6;103(6):829–57.

6. Scuderi S, Altobelli GG, Cimini V, Coppola G, Vaccarino FM. Cell-to-Cell Adhesion and Neurogenesis in Human Cortical Development: A Study Comparing 2D Monolayers with 3D Organoid Cultures. Stem Cell Rep. 2021 Feb 9;16(2):264–80.

7. Vornholt E, Luo D, Qiu W, McMichael GO, Liu Y, Gillespie N, et al. Postmortem brain tissue as an underutilized resource to study the molecular pathology of neuropsychiatric disorders across different ethnic populations. Neurosci Biobehav Rev. 2019 July 1;102:195–207.

8. Kshirsagar A, Mnatsakanyan H, Kulkarni S, Guo J, Cheng K, Ofria LD, et al. Multi-Region Brain Organoids Integrating Cerebral, Mid-Hindbrain, and Endothelial Systems. Adv Sci. 2025;12(33):e03768.

9. Szutorisz H, Egervari G, Sperry J, Carter JM, Hurd YL. Cross-generational THC exposure alters the developmental sensitivity of ventral and dorsal striatal gene expression in male and female offspring. Neurotoxicol Teratol. 2016;58:107–14.

10. Guennewig B, Bitar M, Obiorah I, Hanks J, O’Brien EA, Kaczorowski DC, et al. THC exposure of human iPSC neurons impacts genes associated with neuropsychiatric disorders. Transl Psychiatry. 2018 Apr 25;8(1):89.

11. Andrews, S. (2010). FastQC: A Quality Control Tool for High Throughput Sequence Data [Online]. Available online at: [http://www.bioinformatics.babraham.ac.uk/projects/fastqc/](http://www.bioinformatics.babraham.ac.uk/projects/fastqc/).

12. Ewels P, Magnusson M, Lundin S, Käller M. MultiQC: summarize analysis results for multiple tools and samples in a single report. Bioinformatics. 2016 Oct 1;32(19):3047–8.

13. Krueger F. Trim Galore: a wrapper tool around Cutadapt and FastQC to consistently apply quality and adapter trimming to FastQ files, 2020. [https://www.bioinformatics.babraham.ac.uk/projects/trim_galore/](https://www.bioinformatics.babraham.ac.uk/projects/trim_galore/). Accessed 4 Sept 2025.

14. Dobin A, Davis CA, Schlesinger F, Drenkow J, Zaleski C, Jha S, et al. STAR: ultrafast universal RNA-seq aligner. Bioinforma Oxf Engl. 2013 Jan 1;29(1):15–21.

15. Patro R, Duggal G, Love MI, Irizarry RA, Kingsford C. Salmon provides fast and bias-aware quantification of transcript expression. Nat Methods. 2017 Apr;14(4):417–9.

16. Soneson C, Love MI, Robinson MD. Differential analyses for RNA-seq: transcript-level estimates improve gene-level inferences [Internet]. F1000Research; 2016 [cited 2025 Nov 21]. Available from: https://f1000research.com/articles/4-1521

17. Love MI, Huber W, Anders S. Moderated estimation of fold change and dispersion for RNA-seq data with DESeq2. Genome Biol. 2014;15(12):550.

18. R Core Team (2025). _R: A Language and Environment for Statistical Computing_. R Foundation for Statistical Computing, Vienna, Austria. <[https://www.R-project.org/](https://www.r-project.org/)\>.

19. Xu S, Hu E, Cai Y, Xie Z, Luo X, Zhan L, et al. Using clusterProfiler to characterize multiomics data. Nat Protoc. 2024 Nov;19(11):3292–320.

20. Banerjee-Basu S, Packer A. SFARI Gene: an evolving database for the autism research community. Dis Model Mech. 2010 Mar 8;3(3–4):133–5.

21. Eagles NJ, Wilton R, Jaffe AE, Collado-Torres L. BiocMAP: a Bioconductor-friendly, GPU-accelerated pipeline for bisulfite-sequencing data. BMC Bioinformatics. 2023 Sept 13;24(1):340.

22. Wilton R, Budavari T, Langmead B, Wheelan SJ, Salzberg SL, Szalay AS. Arioc: high-throughput read alignment with GPU-accelerated exploration of the seed-and-extend search space. PeerJ. 2015;3:e808.

23. Krueger F, Andrews SR. Bismark: a flexible aligner and methylation caller for Bisulfite-Seq applications. Bioinforma Oxf Engl. 2011 June 1;27(11):1571–2.

24. Hansen KD, Langmead B, Irizarry RA. BSmooth: from whole genome bisulfite sequencing reads to differentially methylated regions. Genome Biol. 2012 Oct 3;13(10):R83.

25. LeRoith D, Holly JMP, Forbes BE. Insulin-like growth factors: Ligands, binding proteins, and receptors. Mol Metab. 2021 Oct 1;52:101245.

26. Koch S, Claesson-Welsh L. Signal Transduction by Vascular Endothelial Growth Factor Receptors. Cold Spring Harb Perspect Med. 2012 July;2(7):a006502.

27. Blobe GC, Schiemann WP, Lodish HF. Role of Transforming Growth Factor β in Human Disease. N Engl J Med. 2000 May 4;342(18):1350–8.

28. Liu H, Meng L bing, Liu Q. Role of Col1a2 and Postn in left ventricular noncompaction cardiomyopathy. J Cardiothorac Surg. 2025 July 5;20:287.

29. Kapustin AN, Tsakali SS, Whitehead M, Chennell G, Wu MY, Molenaar C, et al. Matrix-associated extracellular vesicles modulate human smooth muscle cell adhesion and directionality by presenting collagen VI. Cooper JA, editor. eLife. 2025 Sept 30;12:RP90375.

30. Mohammadzadeh N, Lunde IG, Andenæs K, Strand ME, Aronsen JM, Skrbic B, et al. The extracellular matrix proteoglycan lumican improves survival and counteracts cardiac dilatation and failure in mice subjected to pressure overload. Sci Rep. 2019 June 24;9(1):9206.

31. Gopal S, Multhaupt HAB, Pocock R, Couchman JR. Cell-extracellular matrix and cell-cell adhesion are linked by syndecan-4. Matrix Biol J Int Soc Matrix Biol. 2017 July;60–61:57–69.

32. Myo Min KK, Ffrench CB, McClure BJ, Ortiz M, Dorward EL, Samuel MS, et al. Desmoglein-2 as a cancer modulator: friend or foe? Front Oncol. 2023 Dec 22;13:1327478.

33. Revollo JR, Grimm AA, Imai S ichiro. The regulation of nicotinamide adenine dinucleotide biosynthesis by Nampt/PBEF/visfatin in mammals. Curr Opin Gastroenterol. 2007 Mar;23(2):164.

34. Berger P, Sirkowski EE, Scherer SS, Suter U. Expression analysis of the N-Myc downstream-regulated gene 1 indicates that myelinating Schwann cells are the primary disease target in hereditary motor and sensory neuropathy-Lom. Neurobiol Dis. 2004 Nov;17(2):290–9.

35. Zhang P, Tchou-Wong KM, Costa M. Egr-1 mediates hypoxia-inducible transcription of the NDRG1 gene through an overlapping Egr-1/Sp1 binding site in the promoter. Cancer Res. 2007 Oct 1;67(19):9125–33.

36. Cabibbo A, Pagani M, Fabbri M, Rocchi M, Farmery MR, Bulleid NJ, et al. ERO1-L, a human protein that favors disulfide bond formation in the endoplasmic reticulum. J Biol Chem. 2000 Feb 18;275(7):4827–33.

37. Acosta-Iborra B, Gil-Acero AI, Sanz-Gómez M, Berrouayel Y, Puente-Santamaría L, Alieva M, et al. Bhlhe40 Regulates Proliferation and Angiogenesis in Mouse Embryoid Bodies under Hypoxia. Int J Mol Sci. 2024 July 12;25(14):7669.

38. Baier PC, Brzózka MM, Shahmoradi A, Reinecke L, Kroos C, Wichert SP, et al. Mice Lacking the Circadian Modulators SHARP1 and SHARP2 Display Altered Sleep and Mixed State Endophenotypes of Psychiatric Disorders. PLOS ONE. 2014 Oct 23;9(10):e110310.

39. Carballo E, Lai WS, Blackshear PJ. Feedback Inhibition of Macrophage Tumor Necrosis Factor-α Production by Tristetraprolin. Science. 1998 Aug 14;281(5379):1001–5.

40. Nicole O, Docagne F, Ali C, Margaill I, Carmeliet P, MacKenzie ET, et al. The proteolytic activity of tissue-plasminogen activator enhances NMDA receptor-mediated signaling. Nat Med. 2001 Jan;7(1):59–64.

41. Dupuis JP, Nicole O, Groc L. NMDA receptor functions in health and disease: Old actor, new dimensions. Neuron. 2023 Aug 2;111(15):2312–28.

42. Kumar KG, Trevaskis JL, Lam DD, Sutton GM, Koza RA, Chouljenko VN, et al. Identification of Adropin as a Secreted Factor Linking Dietary Macronutrient Intake with Energy Homeostasis and Lipid Metabolism. Cell Metab. 2008 Dec 6;8(6):468–81.

43. Banerjee S, Ghoshal S, Girardet C, DeMars KM, Yang C, Niehoff ML, et al. Adropin correlates with aging-related neuropathology in humans and improves cognitive function in aging mice. Npj Aging Mech Dis. 2021 Aug 30;7(1):23.

44. Paul V, Tonchev AB, Henningfeld KA, Pavlakis E, Rust B, Pieler T, et al. Scratch2 Modulates Neurogenesis and Cell Migration Through Antagonism of bHLH Proteins in the Developing Neocortex. Cereb Cortex. 2014 Mar 1;24(3):754–72.

45. Wight TN, Kang I, Evanko SP, Harten IA, Chang MY, Pearce OMT, et al. Versican—A Critical Extracellular Matrix Regulator of Immunity and Inflammation. Front Immunol. 2020 Mar 24;11:512.

46. Harrison OJ, Brasch J, Lasso G, Katsamba PS, Ahlsen G, Honig B, et al. Structural basis of adhesive binding by desmocollins and desmogleins. Proc Natl Acad Sci. 2016 June 28;113(26):7160–5.

47. Peng W, Tan C, Mo L, Jiang J, Zhou W, Du J, et al. Glucose transporter 3 in neuronal glucose metabolism: Health and diseases. Metabolism. 2021 Oct;123:154869.

48. Gao C, Frausto SF, Guedea AL, Tronson NC, Jovasevic V, Leaderbrand K, et al. IQGAP1 Regulates NR2A Signaling, Spine Density, and Cognitive Processes. J Neurosci. 2011 June 8;31(23):8533–42.

49. Su Y, Wang Y, Zhou Y, Zhu Z, Zhang Q, Zhang X, et al. Macrophage migration inhibitory factor activates inflammatory responses of astrocytes through interaction with CD74 receptor. Oncotarget. 2016 Dec 1;8(2):2719–30.

50. Prevot V, Hanchate NK, Bellefontaine N, Sharif A, Parkash J, Estrella C, et al. Function-related structural plasticity of the GnRH system: A role for neuronal–glial–endothelial interactions. Front Neuroendocrinol. 2010 July 1;31(3):241–58.

51. Dautant A, Meier T, Hahn A, Tribouillard-Tanvier D, di Rago JP, Kucharczyk R. ATP Synthase Diseases of Mitochondrial Genetic Origin. Front Physiol [Internet]. 2018 Apr 4 [cited 2025 Dec 1];9. Available from: https://www.frontiersin.org/journals/physiology/articles/10.3389/fphys.2018.00329/full

52. Aïd S, Bosetti F. Targeting cyclooxygenases-1 and −2 in neuroinflammation: therapeutic implications. Biochimie. 2011 Jan;93(1):46–51.

53. Cazzalini O, Scovassi AI, Savio M, Stivala LA, Prosperi E. Multiple roles of the cell cycle inhibitor p21CDKN1A in the DNA damage response. Mutat Res Mutat Res. 2010 Apr 1;704(1):12–20.

54. Yuan A, Sasaki T, Kumar A, Peterhoff CM, Rao MV, Liem RK, et al. Peripherin Is a Subunit of Peripheral Nerve Neurofilaments: Implications for Differential Vulnerability of CNS and Peripheral Nervous System Axons. J Neurosci. 2012 June 20;32(25):8501–8.

55. Roome RB, Bourojeni FB, Mona B, Rastegar-Pouyani S, Blain R, Dumouchel A, et al. Phox2a Defines a Developmental Origin of the Anterolateral System in Mice and Humans. Cell Rep. 2020 Nov 24;33(8):108425.

56. Odell ID, Steach H, Gauld SB, Reinke-Breen L, Karman J, Carr TL, et al. Epiregulin is a dendritic cell-derived EGFR ligand that maintains skin and lung fibrosis. Sci Immunol. 2022 Dec 16;7(78):eabq6691.

57. Crux NB, Elahi S. Human Leukocyte Antigen (HLA) and Immune Regulation: How Do Classical and Non-Classical HLA Alleles Modulate Immune Response to Human Immunodeficiency Virus and Hepatitis C Virus Infections? Front Immunol. 2017 July 18;8:832.

58. Angerer H, Radermacher M, Mańkowska M, Steger M, Zwicker K, Heide H, et al. The LYR protein subunit NB4M/NDUFA6 of mitochondrial complex I anchors an acyl carrier protein and is essential for catalytic activity. Proc Natl Acad Sci U S A. 2014 Apr 8;111(14):5207–12.

59. Lee S, Walker CL, Karten B, Kuny SL, Tennese AA, O’Neill MA, et al. Essential role for the Prader-Willi syndrome protein necdin in axonal outgrowth. Hum Mol Genet. 2005 Mar 1;14(5):627–37.

60. Jay P, Rougeulle C, Massacrier A, Moncla A, Mattel MG, Malzac P, et al. The human necdin gene, NDN, is maternally imprinted and located in the Prader-Willi syndrome chromosomal region. Nat Genet. 1997 Nov;17(3):357–61.

61. Hidalgo J, Aschner M, Zatta P, Vašák M. Roles of the metallothionein family of proteins in the central nervous system. Brain Res Bull. 2001 May 15;55(2):133–45.

62. Zou Y, Donkervoort S, Salo AM, Foley AR, Barnes AM, Hu Y, et al. P4HA1 mutations cause a unique congenital disorder of connective tissue involving tendon, bone, muscle and the eye. Hum Mol Genet. 2017 June 15;26(12):2207–17.

63. Powers RM, Hevner RF, Halpain S. The Neuron Navigators: Structure, function, and evolutionary history. Front Mol Neurosci. 2023 Jan 12;15:1099554.

64. Nobles RD, Zhang C, Müller U, Betz H, McCall MA. Selective Glycine Receptor α2 Subunit Control of Crossover Inhibition between the On and Off Retinal Pathways. J Neurosci. 2012 Mar 7;32(10):3321–32.

65. Hu Y, Cao K, Wang F, Wu W, Mai W, Qiu L, et al. Dual roles of hexokinase 2 in shaping microglial function by gating glycolytic flux and mitochondrial activity. Nat Metab. 2022 Dec;4(12):1756–74.

66. Cho K. Emerging Roles of Complement Protein C1q in Neurodegeneration. Aging Dis. 2019 June 1;10(3):652–63.

67. Müller M, Mazur AJ, Behrmann E, Diensthuber RP, Radke MB, Qu Z, et al. Functional characterization of the human α-cardiac actin mutations Y166C and M305L involved in hypertrophic cardiomyopathy. Cell Mol Life Sci CMLS. 2012 May 29;69(20):3457–79.

68. Tharp CA, Haywood ME, Sbaizero O, Taylor MRG, Mestroni L. The Giant Protein Titin’s Role in Cardiomyopathy: Genetic, Transcriptional, and Post-translational Modifications of TTN and Their Contribution to Cardiac Disease. Front Physiol. 2019 Nov 28;10:1436.

69. Clevers H. Wnt/β-Catenin Signaling in Development and Disease. Cell. 2006 Nov 3;127(3):469–80.

70. Dini G, Verrotti A, Gorello P, Soliani L, Cordelli DM, Antona V, et al. NFIA haploinsufficiency: case series and literature review. Front Pediatr [Internet]. 2023 Oct 17 [cited 2025 Dec 1];11. Available from: https://www.frontiersin.org/journals/pediatrics/articles/10.3389/fped.2023.1292654/full

71. Busquets-Garcia A, Oliveira da Cruz JF, Terral G, Pagano Zottola AC, Soria-Gómez E, Contini A, et al. Hippocampal CB1 Receptors Control Incidental Associations. Neuron. 2018 Sept 19;99(6):1247–1259.e7.

72. Castillo PE, Younts TJ, Chávez AE, Hashimotodani Y. Endocannabinoid signaling and synaptic function. Neuron. 2012 Oct 4;76(1):70–81.

73. Bara A, Ferland JMN, Rompala G, Szutorisz H, Hurd YL. Cannabis and synaptic reprogramming of the developing brain. Nat Rev Neurosci. 2021 July;22(7):423–38.

74. Frischknecht R, Gundelfinger ED. The brain’s extracellular matrix and its role in synaptic plasticity. Adv Exp Med Biol. 2012;970:153–71.

75. Inestrosa NC, Varela-Nallar L. Wnt signaling in the nervous system and in Alzheimer’s disease. J Mol Cell Biol. 2014 Feb 1;6(1):64–74.

76. Hebert-Chatelain E, Desprez T, Serrat R, Bellocchio L, Soria-Gomez E, Busquets-Garcia A, et al. A cannabinoid link between mitochondria and memory. Nature. 2016 Nov 24;539(7630):555–9.

77. Rubino T, Parolaro D. The Impact of Exposure to Cannabinoids in Adolescence: Insights From Animal Models. Biol Psychiatry. 2016 Apr 1;79(7):578–85.

78. Dityatev A, Schachner M, Sonderegger P. The dual role of the extracellular matrix in synaptic plasticity and homeostasis. Nat Rev Neurosci. 2010 Nov;11(11):735–46.

79. Piper M, Barry G, Hawkins J, Mason S, Lindwall C, Little E, et al. NFIA Controls Telencephalic Progenitor Cell Differentiation through Repression of the Notch Effector Hes1. J Neurosci. 2010 July 7;30(27):9127–39.

80. Lipstein N, Verhoeven-Duif NM, Michelassi FE, Calloway N, Hasselt PM van, Pienkowska K, et al. Synaptic UNC13A protein variant causes increased neurotransmission and dyskinetic movement disorder. J Clin Invest. 2017 Mar 1;127(3):1005–18.

